# Rerouting of an Esx substrate pair from the ESX-1 type VII secretion system to ESX-5 by modifying a PE/PPE substrate pair

**DOI:** 10.1101/2020.01.15.906685

**Authors:** Merel P.M. Damen, Trang H. Phan, Roy Ummels, Alba Rubio-Canalejas, Wilbert Bitter, Edith N.G. Houben

## Abstract

Type VII secretion systems (T7SSs) secrete a wide range of extracellular proteins that play important roles in bacterial viability and in host-pathogen interactions of pathogenic mycobacteria. There are five subtypes of mycobacterial T7SSs, called ESX-1 to ESX-5, and four classes of T7SS substrates, namely the Esx, PE, PPE and Esp proteins. At least some of these substrates are secreted as heterodimers. The ESX systems mediate the secretion of specific members of the Esx, PE and PPE proteins, raising the question how these substrates are recognized in a system-specific fashion. PE/PPE heterodimers interact with their cognate EspG chaperones, which recently has been shown to determine their designated secretion pathway. Both structural and pulldown analysis suggest that EspG is unable to interact with Esx proteins and therefore the determining factor for system-specificity of these substrates remains unknown. In this study, we have investigated the secretion specificity of the ESX-1 substrate pair EsxB_1/EsxA_1 (MMAR_0187/MMAR _0188) in *Mycobacterium marinum*. While this substrate pair was hardly secreted when ectopically expressed, secretion was observed when EsxB_1/EsxA_1 was co-expressed together with PE35/PPE68_1 (MMAR_0185/MMAR_0186), which are encoded by the same operon. Surprisingly, co-expressing EsxB_1/EsxA_1 with a modified PE35/PPE68_1 version that carried the EspG_5_ chaperone binding domain, previously shown to redirect this substrate pair to the ESX-5 system, also resulted in co-secretion of EsxB_1/EsxA_1 via ESX-5. Our data suggest a secretion model in which PE35/PPE68_1 is a determinant factor for the system-specific secretion of EsxB_1/EsxA_1.

## Introduction

Mycobacteria possess an unusual hydrophobic cell envelope that protects them from various stresses and contributes to the resilience of pathogenic mycobacteria during infection. Classified as high-GC Gram-positive bacteria, the cell envelope of mycobacteria consists of a standard cell membrane with a surrounding peptidoglycan layer. However, mycobacteria belong to a subgroup of high-GC Gram-positive bacteria that have acquired an extra hydrophobic layer of long-chain fatty acids, called mycolic acids. These specific lipids are covalently linked via an arabinogalactan layer to the peptidoglycan layer, forming a highly rigid and impermeable structure. Mycobacteria employ specialized machineries, called type VII secretion systems (T7SSs) to secrete proteins across their complex cell envelope (1, 2). *Mycobacterium tuberculosis* possesses five of such T7SSs, named ESX-1 to ESX-5 (1, 2), of which ESX-1, ESX-3 and ESX-5 have been functionally analysed (3–10). Each of these systems plays a different role in the mycobacterial life cycle. For example, ESX-1 has a key role in virulence of pathogenic mycobacteria, as it mediates phagosomal rupture inside macrophages (11–14) and the subsequent escape of *M. tuberculosis* from the phagolysosome (3–6, 15, 16). ESX-3 and ESX-5 are necessary for iron and fatty acid uptake, respectively, making them essential for bacterial viability (7–10). Besides their roles in nutrient and metabolite acquisition, ESX-3 and ESX-5 have also been shown to be involved in immune modulation of the host (9, 17, 18).

The substrates that are secreted by these three ESX systems belong to distinctive protein families, *i.e.* Esx, PE, PPE and Esp proteins, most of them belonging to the so-called EsxAB clan protein superfamily (Pfam CL0352) (19). Some of these substrates have been shown to form heterodimers in the cytosol, *i.e.* two Esx proteins pair together and PE proteins pair with a PPE protein, and are thought to be secreted as (partially) folded dimers (13, 20–23). Crystal structures have been solved for several heterodimeric substrates of different ESX systems, revealing highly conserved features, in which the interface of Esx heterodimers (24, 25) as well as the interface of PE/PPE heterodimers, is formed by two pairs of alpha-helices oriented antiparallel to each other (21, 22, 26). Interestingly, each ESX system secretes its own subset of Esx, PE and PPE substrates that are most-likely responsible for the various roles of ESX systems in the bacterial life cycle. How these structurally similar proteins are specifically targeted to their corresponding ESX system still remains unclear. A conserved secretion signal (YxxxD/E) was identified, which is located directly after the helix-turn-helix domain of one partner protein of the Esx heterodimer and the PE partner of the PE/PPE heterodimer. This signal, although required for secretion, was shown to be exchangeable among PE substrates of different ESX systems without changing their initial secretion route (27, 28). Hence, this signal does not determine system-specific secretion of these T7SS substrates.

Structural analysis showed that PPE proteins have a relatively hydrophobic helical tip domain that extends from the characteristic four-helix bundle formed by the PE-PPE interface (21, 22). This helical tip domain is recognized by a cytosolic chaperone, called EspG, in a system-specific manner and this interaction is required for secretion of the PE/PPE pair (21, 22, 29). Subsequently, we could establish the redirection of the ESX-1 substrate pair PE35/PPE68_1 to the ESX-5 system by replacing the EspG_1_ chaperone binding domain with the equivalent domain of the ESX-5 substrate PPE18, suggesting this domain determines through which system these substrates are transported (30). The remaining question is how the Esx substrate pairs that lack this extended tip domain, are specifically recognized and targeted to their designated systems.

Here, we investigated the signals that determine the system-specificity of Esx substrates in *M. marinum* using the ESX-1 heterodimer EsxB_1/EsxA_1 as model substrates. Its encoding genes *esxB_1/esxA_1* (MMAR_0187/MMAR_0188) are adjacent to *pe35/ppe68_1* (MMAR_0185/MMAR_0186), which gene products we previously used as a model ESX-1 dependent PE/PPE heterodimer (29, 30). We found that EsxB_1/EsxA_1 secretion via the ESX-1 system is severely enhanced by the co-expression and secretion of PE35/PPE68_1. Surprisingly, we were able to reroute the EsxB_1/EsxA_1 pair to the ESX-5 system by solely exchanging the EspG binding domain in PPE68_1, showing that the PE/PPE pair determines the system-specificity of this Esx pair.

## Results

### EsxB_1/EsxA_1 require co-expression of PE35/PPE68_1 for efficient ESX-1 dependent secretion

To investigate how the system-specific secretion of Esx substrates is achieved, we investigated the secretion of EsxB_1/EsxA_1 in *M. marinum*. The corresponding coding genes (*MMAR_0187/MMAR_0188*) lie adjacent to the gene pair *pe35/ppe68_1* (*MMAR_0185/MMAR_0186*) and are paralogues of the *pe35-ppe68-esxB-esxA* gene cluster located in the *esx-1* locus (Fig. 1A). We introduced a shuttle plasmid containing *esxB_1/esxA_1*, expressed under the constitutive *hsp60* promoter, in WT *M. marinum* (31). We also included WT *M. marinum* containing the previously analyzed *pe35/ppe68_1* gene pair controlled by the same promoter on an integrative plasmid as an ESX-1 substrate control (30). Secretion was analyzed by immunoblotting using the introduced HA and FLAG epitopes at the C-termini of EsxA_1 and PPE68_1, respectively (Fig. 2A). The cytosolic protein GroEL2 was not detected in the culture supernatant fractions of all analyzed cultures, confirming the integrity of the bacterial cells. Consistent with published data (27, 30), PPE68_1-FLAG was detected in the culture supernatant fraction of the WT strain as a faint smeary band of ∼40 kDa (Fig. 2A, lane 14-15). Whereas EsxA_1-HA was well-expressed, as judged from the HA signals detected in the pellet fractions (Fig. 2A, lane 4-5), this protein was not secreted by the WT strain (Fig. 2A, lane 16-17). Surprisingly, secretion of endogenous EsxA, detected by an EsxA antibody, was severely affected by the overexpression of EsxB_1/EsxA_1 (Fig. 2A, lane 16-17). To investigate the impact of EsxB_1/EsxA_1 overexpression also on other ESX-1 substrates, secretion of EspE was investigated (Fig. 3). This substrate remains attached to the bacterial surface in *M. marinum* and can be extracted by the mild detergent Genapol X-080. The presence of EspE in the Genapol extracted fraction was completely abolished in the strain overexpressing EsxB_1/EsxA_1, indicating that this ectopically expressed pair has a broad effect on ESX-1 secretion.

**Figure 1.**
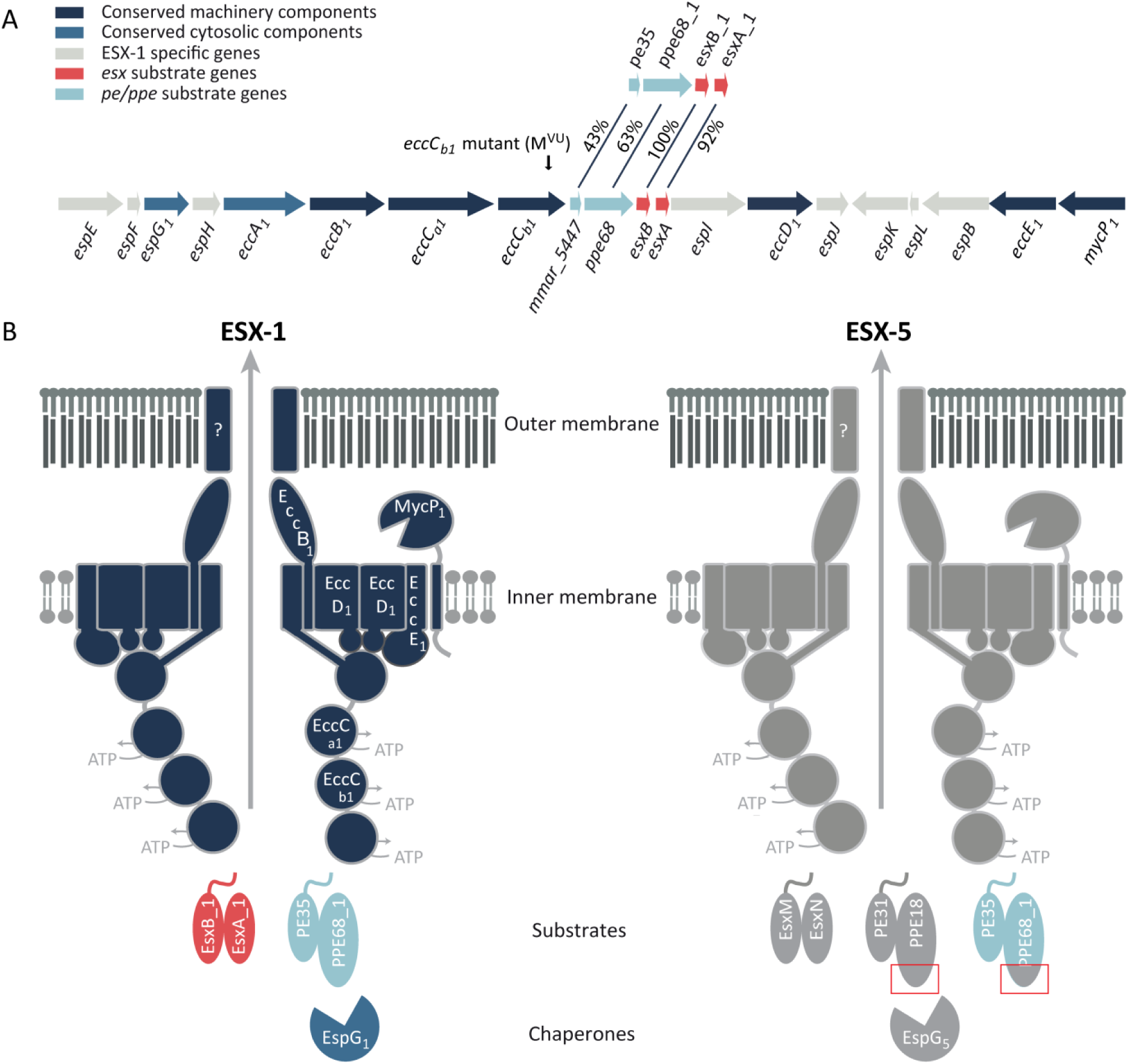
Schematic representation of the *esx-1* locus and type VII secretion systems. (A) Genetic organization of the *esx-1* locus and the duplicated region containing *pe35/ppe68_1/esxB_1/esxA_1*. Genes are color-coded according to the localization of their encoded proteins (see color key). The frameshift mutation in the *eccC*_*b1*_ mutant (also named M^VU^) used in this study is indicated by an arrow. (B) Current model of substrate recognition by the type VII secretion systems. EspG components recognize cognate PE/PPE substrates, independently of the general secretion signal located in the C terminus of the PE proteins. This recognition is required for secretion via the cognate secretion machinery located in the mycobacterial inner membrane. Shown are the model substrates used in this study, *i.e.* the ESX-1 substrate pairs EsxB_1/EsxA_1 and PE35_1/PPE68_1, and the ESX-5 substrate pairs EsxM/EsxN and PE31/PPE18. Previously, PPE68_1 was shown to be rerouted to the ESX-5 system by exchanging the EspG_1_ binding domain with the equivalent domain of PPE18, indicated by red boxes (30).

**Figure 2.**
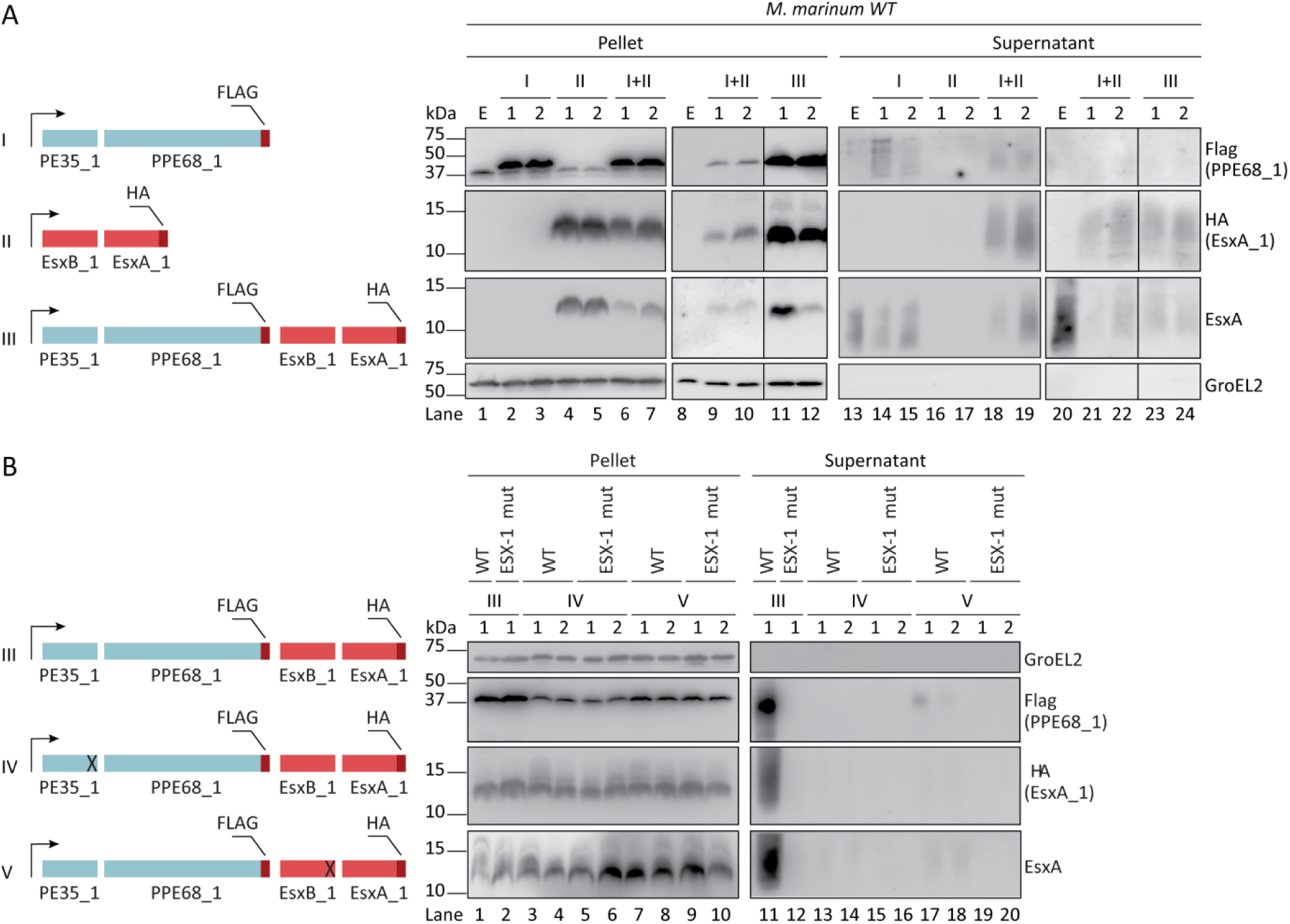
Heterologous EsxA_1/EsxB_1 require co-expression and secretion of PE35/PPE68_1 for efficient secretion via the ESX-1 system (A) A schematic representation of the different constructs used are shown on the left. The four genes encoding the *M. marinum* ESX-1 substrates PE35/PPE68_1 (in blue) and EsxB_1/EsxA_1 (in pink) are either expressed from separate *hsp60* promoters (I and II) or co-expressed under the same *hsp60* promoter (III). Immunoblot analysis of the cell pellet and culture supernatant fractions of WT *M. marinum* using an HA antibody to detect EsxA_1-HA, a FLAG-antibody to detect PPE68_1-FLAG, an EsxA antibody, detecting both endogenous and exogenous EsxA paralogues, and a GroEL2 antibody to detect the intracellular control protein GroEL2. (B) A schematic representation of the different constructs used are shown on the left; deletions are indicated by a cross. Immunoblot analysis using the same antibodies as under A to analyze secretion by WT and the ESX-1 mutant. In all blots, equivalent OD units were loaded; 0.2 OD for pellet and 0.5 OD for supernatant fractions. Numbers indicate two independent *M. marinum* colonies carrying the same construct. E, empty strain.

**Figure 3.**
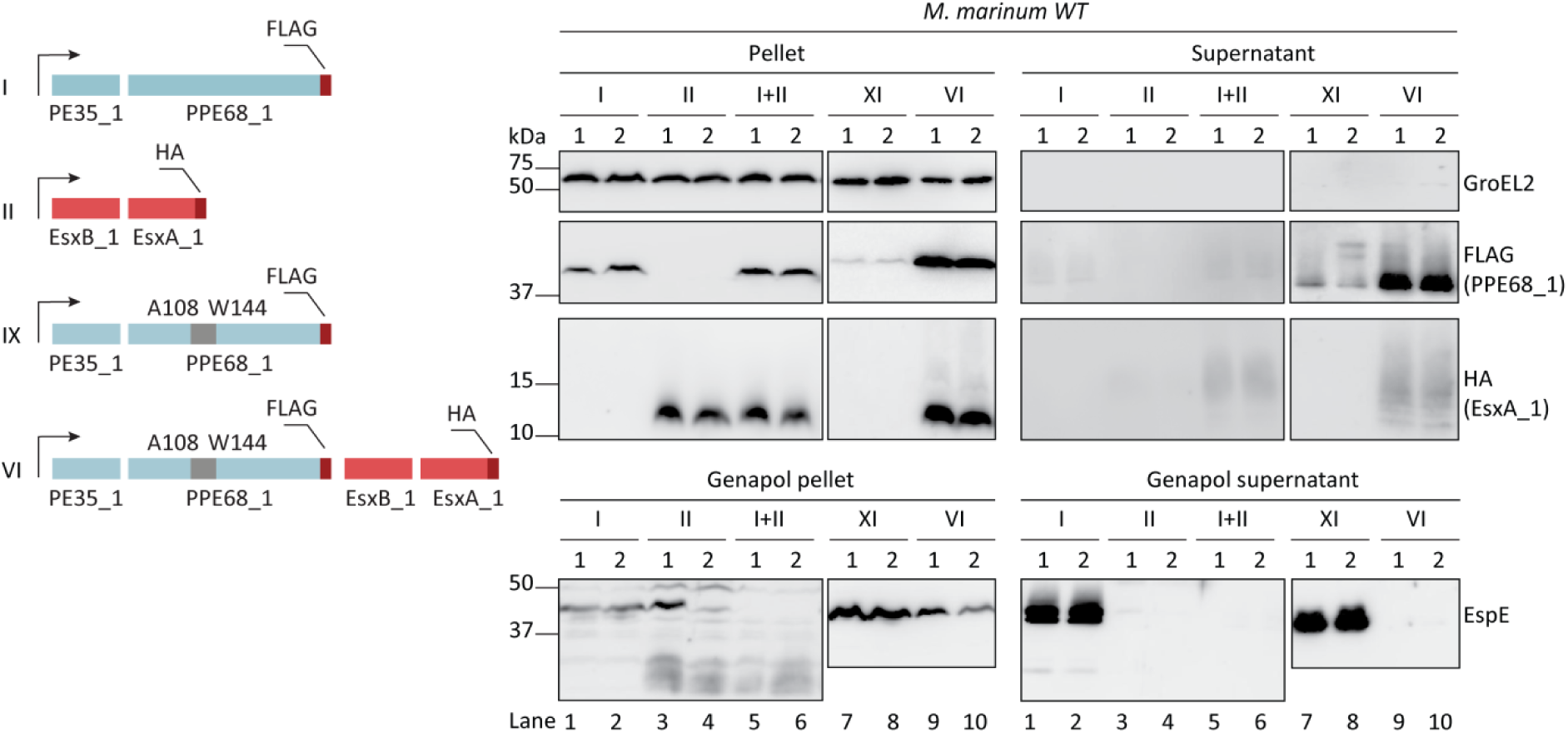
Overexpression of EsxA_1 interferes with EspE secretion and this secretion is not restored by co-overexpressing or rerouting of PE35/PPE68_1 via ESX-5. A schematic representation of the different constructs used are shown on the left. The genes encoding the *M. marinum* ESX-1 substrates PE35/PPE68_1 (in blue), PE35/PPE68_1 containing an EspG_5_ chaperone binding domain (in blue and grey) and EsxB_1/EsxA_1 (in pink) are either expressed from separate *hsp60* promoters (I, II, XI) or co-expressed under the same *hsp60* promoter (VI). Immunoblot analysis of the cell pellet and culture supernatant fractions of WT *M. marinum* using an HA antibody to detect EsxA_1-HA, a FLAG-antibody to detect PPE68_1-FLAG, an EsxA antibody, detecting both endogenous and exogenous EsxA paralogues, and a GroEL2 antibody to detect the intracellular control protein GroEL2. Surface-localized EspE was extracted from bacterial cell pellets using Genapol X-080, resulting in Genapol pellet and supernatant fractions. These fractions were immune-stained with EspE antibodies. Equivalent OD units of cell pellets or Genapol pellet (0.2 OD unit) and culture supernatants or Genapol supernatants (0.5 OD unit) are shown. Numbers indicate two independent *M. marinum* colonies carrying the same construct.

As several T7SS substrates, in particular those of the ESX-1 system, have been shown to be dependent on each other for secretion (30, 32, 33), we hypothesized that the secretion of heterologous EsxB_1/EsxA_1 might require the co-overexpression of the PE35/PPE68_1 pair that is putatively located in the same operon. In addition, similarly organized loci containing a *pe/ppe* pair and an adjacent *esx* gene pair can be observed in other ESX clusters. We co-electroporated the integrative pMV361∷*pe35/ppe68_1-flag* and the pSMT3∷*esxB_1/esxA_1-ha* in WT *M. marinum*. Secretion analysis followed by immunoblotting showed that the co-expression of EsxB_1/EsxA_1-HA did not seem to affect the expression and secretion of PPE68_1-FLAG (Fig. 2A, lane 6-7,9-10 and lane 18-19, 21-22). In contrast, EsxA_1-HA was now efficiently detected as a smeary band of ∼15 kDa in the supernatant of the strain that co-expressed both the PE/PPE and the Esx substrate pairs (Fig. 2A, lane 18-19, 21-22). Furthermore, the immunoblot signals in the supernatant fractions using the EsxA antibody, which probably detects both EsxA_1-HA and endogenous EsxA, became comparable to that of the WT strain without any constructs (Fig. 2A, lane 18-19, 21-22). However, the presence of EspE in the Genapol extracted fraction was not restored in the strain overexpressing both the Esx and PE/PPE pair, suggesting that the secretion of endogenous ESX-1 substrates was still blocked (Fig. 3). These data show that the efficient secretion of overexpressed EsxA_1-HA relies on the co-overexpression of PE35/PPE68_1.

Because the integrative pMV361 plasmid and the multicopy pSMT3 plasmid differ in copy numbers, thereby possibly resulting in suboptimal co-secretion of the two substrate pairs, we also introduced the complete *pe35/ppe68_1/esxB_1/esxA_1* locus into the pSMT3 plasmid again with a FLAG and HA tag fused to the C-termini of PPE68_1 and EsxA_1, respectively. We observed that while the cellular levels of both EsxA_1-HA and PPE68_1-FLAG was increased (Fig. 2A, lane 11-12), the secretion of EsxA_1-HA was similar to the condition when the two substrate pairs were independently expressed (Fig. 2A, lane 23-24). Taken together, our data suggest that the co-expression of PE35/PPE68_1, but not its co-transcription, increases the secretion of ectopically expressed EsxB_1/EsxA_1 in *M. marinum*.

We next investigated whether the heterologous secretion of EsxA_1 requires not only co-overexpression, but also co-secretion of ectopically expressed PPE68_1. To test this, the C-terminal 15 and 21 amino acids of PE35 and EsxB_1, respectively, containing their secretion signals, were deleted, which abolishes their secretion. We analyzed the effect of these truncations on the secretion of PPE68_1 and EsxA_1 in the WT and an *eccCb*_*1*_ mutant strain (ESX-1 mutant), previously described as a non-functional *esx-1* mutant (34) (Fig. 2B). In all tested cultures, the supernatants were devoid of GroEL2, indicating the integrity of the cells (Fig. 2B). Secretion of both EsxA-HA and PPE68_1-FLAG was abolished in the ESX-1 mutant strain, confirming these proteins are secreted via the ESX-1 secretion system (Fig. 2B, lane 12). Importantly, immunoblot analysis showed that the deletion of the C-terminal tail of either PE35 or EsxB_1 disrupted the secretion of both PPE68_1 and EsxA_1 in WT *M. marinum* (Fig. 2B, lane 13-14, 17-18). Only some minor secretion of PPE68_1 was observed in the WT strain, but not in the ESX-1 mutant strain, when the secretion signal in EsxB_1 was deleted (Fig. 2B, lane 17-18). We therefore conclude that the presence of both the PE35 and EsxB_1 C-terminal tails, containing the secretion signals, are required for the secretion of both substrates. This indicates that EsxB_1/EsxA_1 and PE35/PPE68_1 are co-dependently secreted via the ESX-1 system.

### WT EsxB_1/EsxA_1 is rerouted to the ESX-5 system by introducing the EspG_5_ binding domain in PPE68_1

Two different domains in T7SS substrates have been identified that are important for secretion. On the one hand the YxxxD/E motif, located directly after the helix-turn-helix domain of some Esx and all PE proteins, and secondly the EspG binding domain of PPE proteins. Although the C-terminal tails of PE substrates, secreted through different ESX systems, can in general be exchanged without changing their predetermined secretion route (27), we were able to redirect PPE68_1 to the ESX-5 system by exchanging its EspG_1_ chaperone binding domain with the equivalent domain of the ESX-5 substrate PPE18 (30). We therefore tested which of these signals is required to establish rerouting of EsxA_1 to the ESX-5 system. For this, we constructed *pe35/ppe68_1/esxB_1/esxA_1*, in which *ppe68_1* was engineered to express a PPE68_1 variant that carries the EspG_5_ binding domain of the PPE18 protein (Fig. 1B), as previously described (30). This construct was named SINGLE SWAP (Fig. 4A) and the modified PPE68_1 protein was named PPE68_1 SWAP. To additionally investigate the role of the EsxB_1 secretion signal, the C-terminal domain EsxB_1 containing the YxxxD/E motif was replaced by the equivalent region of the ESX-5 substrate EsxM (Fig. 4B; Fig. 1B). This construct, also containing PPE68_1 SWAP was designated DOUBLE SWAP (Fig. 4A). Finally, we also replaced the 15 amino-acid C-terminal domain of the PE35 by the corresponding region of the ESX-5 substrate PE31 (Fig. 1B), resulting in the construct of TRIPLE SWAP (Fig. 4A). The four constructs, *i.e.* WT, SINGLE, DOUBLE and TRIPLE SWAP, were introduced in *M. marinum* WT, the *eccC*_*b1*_ mutant (ESX-1 mutant) and an *ΔeccC*_*5*_ (ESX-5 mutant) strain, after which secretion was analyzed.

**Figure 4.**
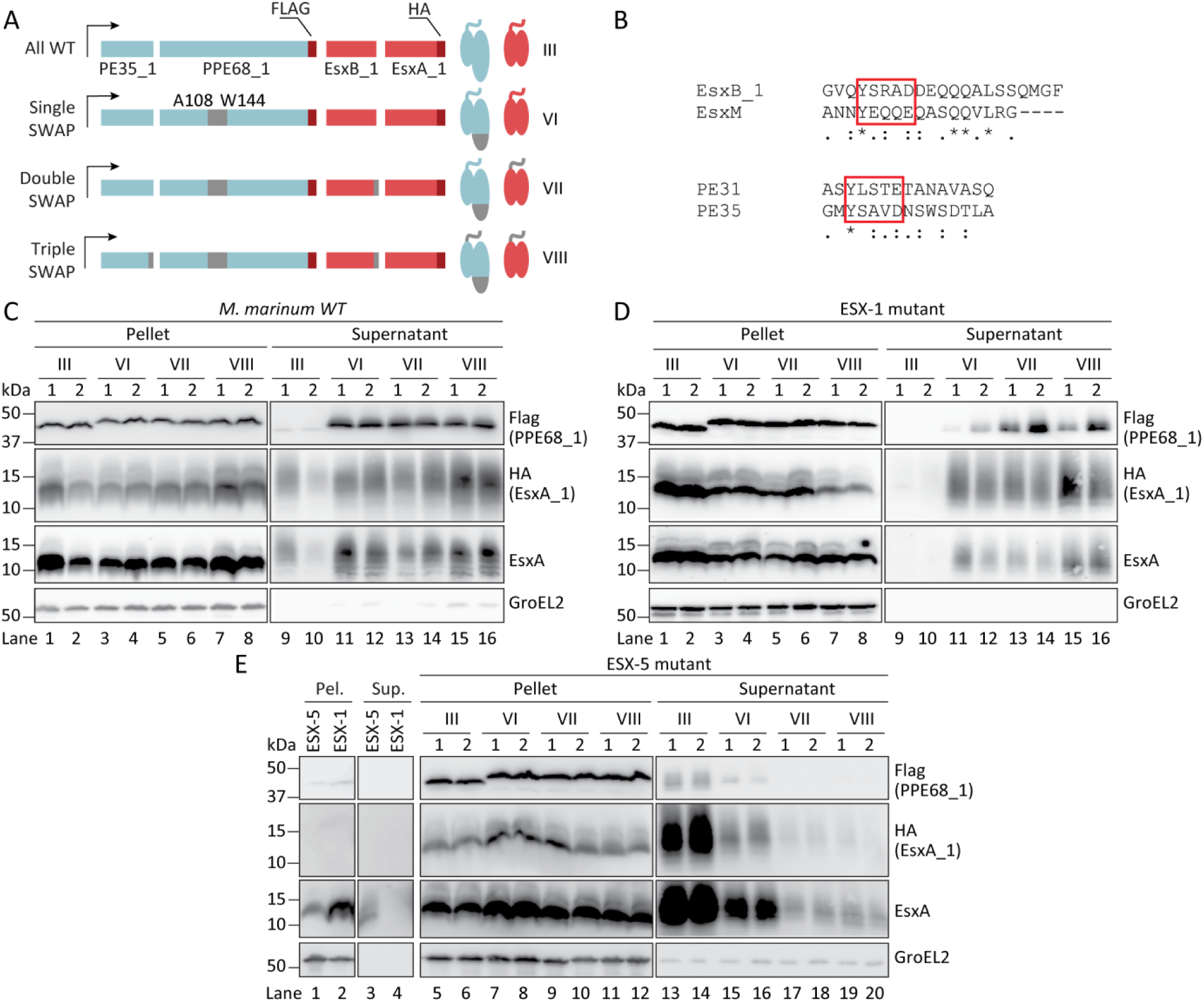
EsxA_1 is co-rerouted to ESX-5 by introducing the EspG_5_ binding domain of ESX-5 substrate PPE18 into PPE68_1. (A) Schematic representation of the different constructs used in the secretion analysis. The introduced sequences of the ESX-5 substrates PE31/PPE18 are in grey. The FLAG-tag (on PPE68_1) and HA-tag (on EsxA_1) are shown in red. (B) Alignment of the swapped sequences containing the C-terminal secretion motifs of the ESX-5 substrates PE31 and EsxM and the ESX-1 substrates PE35 and EsxB_1, respectively. The conserved secretion motif YxxxD/E are highlighted with a red box. (C,D,E) Immunoblot analysis of EsxA_1 as detected with an HA antibody, PPE68_1 as probed with a FLAG antibody, endogenous and exogenous EsxA paralogues using an EsxA antibody and intracellular GroEL2 by an GroEL2 antibody, in pellet and supernatant fractions. Different derivatives of PE35/PPE68_1/EsxB_1/EsxA_1 were tested in WT *M. marinum* (C), an *eccCb*_*1*_ mutant (ESX-1 mutant; D), and an *eccC*_*5*_ knockout strain (ESX-5 mutant; E). Equivalent OD units of cell pellets (0.2 OD unit) and culture supernatants (0.5 OD unit) are shown. Numbers indicate two independent *M. marinum* colonies carrying the same construct.

Interestingly, we observed that the presence of the SINGLE, DOUBLE and TRIPLE SWAP constructs seemed to cause some minor lysis of WT *M. marinum* cells, as a small amount of GroEL2 was consistently detected in the supernatants of these cultures (Fig. 4C). Nevertheless, the detected amount of GroEL2 was comparable among the strains expressing the different constructs, allowing further analysis. As observed before, the WT construct resulted in expression and secretion of both PPE68_1-FLAG and EsxA_1HA (Fig. 4C, lane 1-2 and 9-10). The EsxA antibody was included to confirm the total EsxA expression and secretion (Fig. 4C, lane 1-2 and 9-10). As seen previously for the SINGLE SWAP construct (30), we observed that PPE68_1 SWAP, appearing as a slightly higher band than the PPE68_1 WT, was expressed and efficiently secreted in the WT strain (Fig. 4C, lane 3-4 and 11-12). Notably, while secretion of PPE68_1 SWAP was more efficient than PPE68_1 WT (30), the amount of EsxA_1 was also higher in the supernatant fractions, as judged by an increased intensity of both HA and EsxA signals (Fig. 4C, lane 11-12). The presence of the C-terminal tail of EsxM in the DOUBLE SWAP construct did not affect the secretion of both PPE68_1 SWAP and EsxA_1 as similar intensities of the detected signals were observed (Fig. 4C, lane 13-14). However, when ESX-5 secretion signals were introduced in both PE35 and EsxB_1, the secretion of both the PPE68_1 SWAP and EsxA_1 seemed to reach the highest efficiency.

We subsequently addressed the involved secretion systems by first introducing the different constructs in the ESX-1 mutant strain. In contrast to WT cells, GroEL2 was not detected in the supernatant fractions of this mutant strain, indicating the integrity of the cells in the presence of the constructs (Fig. 4D). As observed previously, secretion of both PPE68_1-FLAG and EsxA_1-HA of the WT construct was abrogated (Fig. 4D, lane 9-10). In contrast, the PPE68_1 SWAP protein was still secreted (Fig. 4D, lane 11-12), confirming our previous observation that the PPE68_1 SWAP was secreted independently from the ESX-1 system (30). Importantly, we also still detected EsxA_1 in the supernatant, using both the HA and the EsxA antibody (Fig. 4D, lane 11-12), suggesting that this ESX-1 substrate is secreted in an ESX-1 independent manner as well. This is highly interesting as both EsxA_1 and EsxB_1 are unmodified in the SINGLE SWAP construct. Similar as for the WT bacteria, the DOUBLE SWAP construct showed comparable levels of EsxA_1 secretion as the SINGLE SWAP constructs (Fig. 4D, lane 13-14), while the secretion of EsxA_1 seemed the most efficient in the presence of both the ESX-5 secretion signals in the TRIPLE SWAP construct (Fig. 4D, lane 15-16). Together, these data showed that PPE68_1 SWAP determines the ESX-1 independent secretion of EsxA_1.

To confirm that PPE68_1 SWAP and EsxA_1 are secreted by the ESX-5 system, we introduced the constructs in the *ΔeccC*_*5*_ strain. EccC_5_ is an essential component of the ESX-5 machinery (35, 36) and deletion of this component blocks ESX-5-dependent secretion (7). In this strain, the presence of all tested constructs consistently caused minor bacterial lysis, as GroEL2 was found in all supernatant fractions (Fig. 4E). As a similar phenotype was observed for the SINGLE, DOUBLE and TRIPLE SWAP constructs in WT background, but not in the ESX-1 mutant strain, the bacterial leakage induced upon ectopic expression of these proteins seemed to be linked to a functional ESX-1 system. With the WT construct, we detected both PPE68_1-FLAG and EsxA_1-HA in the supernatant fractions by using the FLAG and HA antibody, respectively (Fig. 4E, lane 13-14). The PPE68_1 SWAP and EsxA_1 of the SINGLE SWAP were moderately detected in the supernatant (Fig. 4E, lane 15-16), whereas they were no longer detected in the supernatants of bacteria containing either the DOUBLE or the TRIPLE SWAP (Fig. 4E, lane 17-20). In the two latter cases, the signals using the EsxA antibody were detected at comparable levels (Fig. 4E, lane 17-20) and were similar to that of the empty *ΔeccC*_*5*_ strain (Fig. 4E, lane 3). Thus, our data show that the secretion of both proteins became mostly dependent on the ESX-5 system when the EspG_5_ binding domain was introduced. The observed residual secretion of PPE68_1-FLAG and EsxA_1-HA with the SINGLE SWAP construct indicates that a small amount of these substrate pairs is still secreted via ESX-1. Interestingly, we previously showed that secretion of PPE68_1 SWAP was completely blocked in the same ESX-5 mutant in the absence of ectopically expressed EsxB_1/EsxA_1 (30). This indicates that this Esx substrate pair might be able to guide some amount of PPE68_1 SWAP to the ESX-1 system.

Finally, we observed a competitive correlation between the secretion of the rerouted substrates PE35/PPE68_1 SWAP and native substrates of the ESX-5 system, the PE_PGRS proteins. Using the Genapol extraction method to analyse the surface localization of PE_PGRS proteins, we observed a lower amount of these proteins in the Genapol extracted fraction in the ESX-1 mutant strains expressing the four different constructs compared to WT bacteria containing the same constructs (Fig. 5). This suggests that, similar to what was reported previously, the redirection of ESX-1 substrates to the ESX-5 system interferes with the export of endogenous ESX-5 substrates (30).

**Figure 5.**
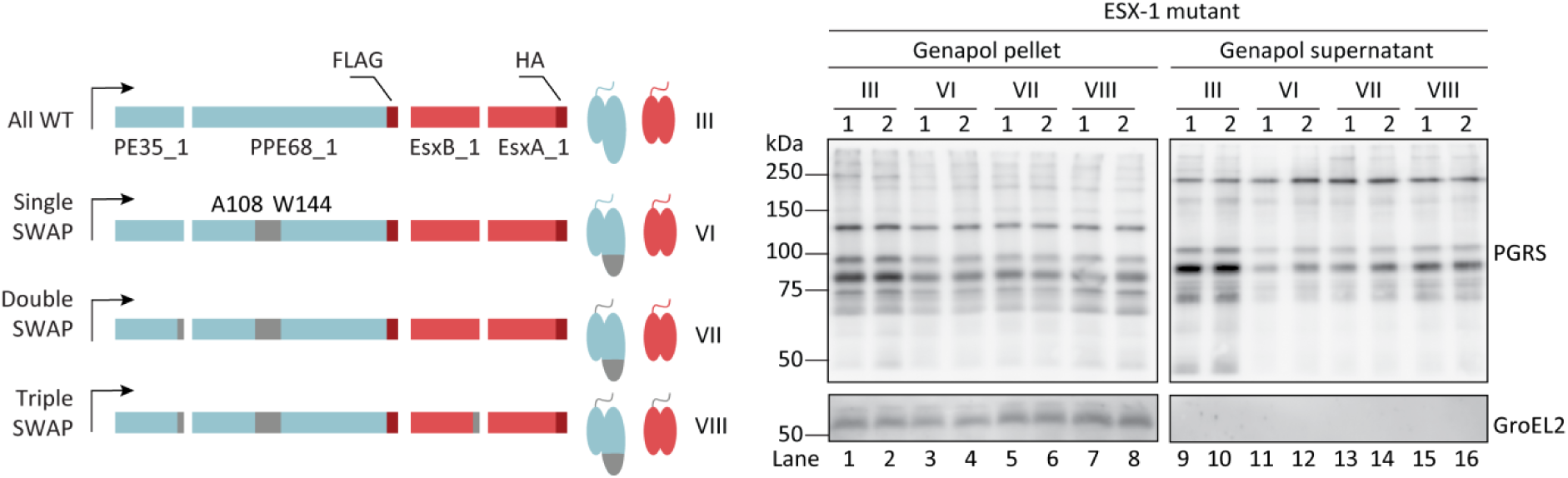
Rerouted PP68_1 and EsxA_1 compete with endogenous ESX-5 substrates for secretion. Immunoblot analysis of the surface-localized ESX-5 dependent PE_PGRS substrates in the ESX-1 mutant strain overexpressing PE35/PPE68_1/EsxB_1/EsxA_1 variants. The fractions of enriched cell surface proteins (Genapol supernatant) were collected from the bacterial cell pellets (Genapol pellet) using Genapol X-080. Equivalent OD units were loaded; 0.2 OD for pellet and 0.5 OD for Genapol supernatants.

In summary, introducing the EspG_5_ binding domain in PPE68_1 resulted in the rerouting of both this PPE substrate and EsxA_1 to the ESX-5 system, not only further confirming that these proteins are co-secreted and but also showing that the PPE protein is involved in determining the system-specificity of the Esx substrate. Introduction of two ESX-5 secretion motifs optimized the secretion efficiency via ESX-5, showing that these signals have system-specific functionality to some extent.

## Discussion

Mycobacterial T7SSs secrete their own subset of Esx, PE and PPE proteins, which resemble each other on sequence and structural levels. This phenomenon raises the question how these substrates are specifically recognized by the T7SS subtypes. Recently, we showed that the system-specificity of the PE/PPE substrates is determined by the EspG chaperone binding domain on the PPE protein (30). However, this cannot apply to the Esx heterodimers, since they lack this chaperone binding domain and therefore cannot interact with the EspG chaperones (37). In this study, we show that the PE35/PPE68_1 heterodimer defines the system-specific secretion of EsxB_1/EsxA_1 in *M. marinum*.

We found that ESX-1 dependent secretion of exogenous EsxB_1/EsxA_1 is severely enhanced when PE35/PPE68_1 were co-expressed. Co-expression of PE35/PPE68_1 from the same promoter was not required for EsxA_1 secretion, suggesting that this dependency is not transcriptionally linked. Our previous observation that deletion of *espG*_*1*_ leads to a loss of not only PE/PPE secretion, but also of EsxB/EsxA and other ESX-1 substrates in *M. marinum* (30, 38), already hinted towards the dependency of EsxB/EsxA on PE/PPE substrates for their secretion. Interestingly, in *M. tuberculosis* the dependency of EsxA secretion on PPE68 seems more complex. Although PPE68 itself is indeed crucial for EsxA secretion, an *espG*_*1*_ deletion in this species, which is predicted to affect PPE68, does not seem to affect EsxA secretion (4, 33). Since there is only a single copy of *esxA* in the genome of *M. tuberculosis*, contrary to the four highly homologous *esxA* copies *M. marinum*, it is possible that the two pathogens employ distinct mechanisms to govern the regulation and/or secretion of EsxA. Alternatively, there might be redundancy of different PE/PPE proteins in *M. tuberculosis* in facilitating the secretion of EsxA, or EspG_1_ is less important for ESX-1 secretion in *M. tuberculosis*.

In this study, we also confirmed that the C-terminal tails of PE35 and EsxB_1 are required for the secretion of the corresponding heterodimer, consistent with other studies (27). Moreover, our findings that these two secretion motifs are strictly required for the secretion of both heterodimers is highly interesting. These data show that the secretion of EsxA_1 is not only dependent on the co-overexpression, but also on the secretion of PE35/PPE68_1. In addition, the observation that secretion of PPE68_1 was diminished when the secretion of EsxB_1/EsxA_1 was blocked by deleting the secretion signal of EsxB_1 suggests that both heterodimers are mutually dependent on each other for their export. Interestingly, the cytosolic accumulation of EsxB_1/EsxA_1, which occurs without co-overexpression of PE35/PPE68_1 or when the C-terminal tails of PE35 and/or EsxB_1 was deleted, also abolishes the secretion of endogenous EsxA and EspE. This is in agreement with our previous observation that the introduction of an ESX-5 secretion signal in PE35 of ectopically expressed PE35/PPE68_1 results in a secretion block of both PPE68_1 and endogenous EsxA (30). Possibly, a cytosol factor that is required for substrate recognition, *e.g.* EspG_1_, is titrated away by these non-secreted substrates.

The finding that WT EsxB_1/EsxA_1 could be rerouted to the ESX-5 system in *M. marinum* by solely manipulating the EspG binding domain of PE35/PPE68_1 is unexpected. The fact that WT EsxB_1/EsxA_1 could be co-rerouted in this manner underlines the strict dependency of EsxB_1/EsxA_1 secretion on PE35/PPE68_1. In addition, secretion of both exogenous PPE68_1 WT via ESX-1 and PPE68_1 SWAP via ESX-5 does not seem to be enhanced upon co-expression of EsxB_1/EsxA_1. This shows there is hierarchy in secretion where the PE/PPE pair controls and might regulate secretion of EsxB_1/EsxA_1. Co-rerouting of endogenous EsxA by the over-expression of PE35/PPE68_1 SWAP was previously not observed (30), suggesting that expression levels of endogenous Esx and PE/PPE pairs are well-balanced. Interestingly, the ability of the ESX-5 system to secrete the two substrate pairs, while they still carry both ESX-1 secretion signals, reflects its flexibility in secreting a wide range of substrates, as already discussed in other studies (21, 22, 34). Nevertheless, rerouting was most efficient when both PE35 and the EsxB_1 carried the ESX-5 C-terminal secretion signals of PE31 and EsxM, respectively. This suggests that the secretion signals of PE31 and EsxM are more optimal for the recognition and secretion by the ESX-5 machinery.

So far, the reason behind co-dependence among T7SS substrates remained unclear. It has been proposed that binding to or activation of the T7SS could explain this phenomenon. Previous studies showed that EsxB binds to the third nucleotide binding domain (NBD) of the conserved membrane complex component EccC_ab1_ and induces hexamerization of this ATPase *in vitro* (28, 39, 40). Specifically, the last seven amino acids of EsxB were shown to be essential for this interaction (28). The C-terminus of other Esx homologs in *M. tuberculosis* are likely to be structured similarly to the C-terminus of EsxB, but do contain different amino acids sequences, suggesting this domain might be involved in system-specific recognition (28, 40, 41). Importantly, structural analysis of EccC of *Thermomonospora curvata* showed that the crucial first NBD is kept in an inactivate state by a specific region in the linker 2 domain that connects the first and second NBD, and binding of EsxB is not able to activate this ATPase activity *in vitro* (40). It was therefore suggested that an additional trigger is necessary to activate EccC, which could be the binding of PE/PPE substrates. From our study, it seems evident that the secretion of the Esx substrates are closely linked to that of PE/PPE heterodimers. Possibly, they bind simultaneously or sequentially in order to activate all three ATPase domains of EccC, after which transport through the membrane complex is achieved. Such a model would explain both the necessity for equal expression levels of both heterodimers, as well as the secretion dependency of the Esx pair on the PE/PPE pair that we observed here. However, PE/PPE proteins can only be found in the genus of *Mycobacterium*, while the homologues of the Esx substrates and the EccC core component can be found in a more diverse repertoire of Gram-positive species (42–44). It will be interesting to see the differences in substrate recognition and secretion between these different systems.

The homologue of EsxA_1, EsxA, is the most-studied T7SS substrate and has been suggested to be responsible for ESX-1-induced phagosomal rupture. EsxA was found to be associated with membrane lysis when a transposon mutant of *esxA/esxB* was unable to lyse cultured lung epithelial cell lines (4, 45). Further genetic studies in *M. marinum* showed that several different transposon mutants defective in EsxA secretion lost haemolytic activity and were attenuated in zebrafish (3, 12, 15, 46), supporting the hypothesis that EsxA is a crucial virulent factor of pathogenic mycobacteria. However, secretion of different ESX-1 substrate classes has been shown to be interdependent on each other (32, 47), *e.g.* loss of EspA or PPE68 secretion led to secretion defects of EsxA and *vice versa* (30, 32). Therefore, studying functions of individual ESX-1 substrates during the mycobacterial infection cycle has been a challenge. While protein sequences of EsxB and EsxB_1 are identical, EsxA_1 shares 92% protein sequence identity with EsxA. Given the high similarity, it has been suggested that EsxB_1/EsxA_1 have an equivalent functionality as the *esx-1* encoded EsxB/EsxA (48). The observation that WT EsxA_1 can be destined for the ESX-5 system provides a unique platform to investigate exact roles of this protein in host-pathogen interactions. Current research is focusing on the redirection of EsxB/EsxA in order to directly assess the membrane lysis activity of this substrate pair.

## Experimental Procedures

### Bacterial strains and growth cultures

All mycobacterial strains were grown on Middlebrook 7H10 plates (Difco) containing OADC supplement (oleic acid, albumin, dextrose and catalase; BD Biosciences) or liquid 7H9 medium containing ADC supplement (BD Biosciences) and the appropriate antibiotics (see below). *M. marinum* strains were grown at 30 °C, 90 rpm. All mycobacterial strains and mutants are listed in Table S1. *Escherichia coli* strain DH5α was used for cloning procedures and plasmid accumulation, and was grown on lysogeny broth (LB) plates or liquid broth at 37 °C, 200 rpm. Growth media was supplemented with the appropriate antibiotics at the following concentrations: kanamycin (Roche) 25 µg/ml; hygromycin (Sigma) 50 µg/ml.

### Plasmid construction

All PCRs were carried out with the Phusion High-Fidelity DNA polymerase (Finnzymes) using primers listed in supplemental Table S2. The restriction sites used for cloning are indicated in this table.

### Protein secretion and immunoblot analysis

*M. marinum* strains were grown in 7H9 liquid medium supplemented with ADC, 0.05% Tween 80, and appropriate antibiotics until mid-logarithmic phase, after which the cells were washed and inoculated in 7H9 medium with 0.2% dextrose, 0.05% Tween 80 at an optical density at 600 nm (OD_600_) of 0.4 and grown for another 16 h. The cells (Pellet) were spun down for 10 min at 6,000 xg, washed with phosphate buffered saline (PBS), and resuspended in SDS loading buffer (containing 100mM DTT and 2% SDS). Supernatants were passed through 0.2 µm-pore-size filter units and proteins were precipitated with trichloroacetic acid (TCA) and resuspended in SDS loading buffer. Alternatively, the cells were resuspended in 0.5% Genapol X-080 and incubated for 1 h at room temperature. Samples were spun down and pellets were resuspended in SDS sample loading buffer (Genapol Pellet), while 5x SDS sample buffer was added to the supernatant containing Genapol X-080 (Genapol Supernatant) to obtain a final concentration of 1x SDS buffer. Proteins were separated on SDS-PAGE gels and transferred to a nitrocellulose membrane, and membranes were stained with anti-GroEL2 (monoclonal antibody Cs44; John Belisle, NIH, Bethesda, MD, USA), anti-PE_PGRS (7C4.1F7) (34), anti-ESAT-6 (monoclonal antibody [MAb] Hyb76-8) (49), anti-HA (HA.11; Covance), anti-EspE (polyclonal rabbit antibody; Eric Brown; Genentech), anti-Flag (M2 monoclonal antibody produced in mouse, Sigma).

## Acknowledgements

We thank Joen Luirink for valuable discussions.

## FOOTNOTES

Funding was provided by a VIDI grant (864.12.006; to THP and ENGH) and an ALW Open grant (ALWOP.319; to MPMD) both from the Netherlands Organization of Scientific Research. The funders had no role in study design, data collection and analysis, decision to publish, or preparation of the manuscript.

## The abbreviations used are

T7SS: Type VII secretion system;
NBD: nucleotide binding domain;
PBS: phosphate buffered saline;
TCA: trichloroacetic acid

## References

1. Houben, E. N. G., Korotkov, K. V, and Bitter, W. (2014) Take five - Type VII secretion systems of mycobacteria. Biochim. Biophys. Acta - Mol. Cell Res. 1843, 1707–1716

2. Bitter, W., Houben, E. N. G., Bottai, D., Brodin, P., Brown, E. J., Cox, J. S., Derbyshire, K., Fortune, S. M., Gao, L.-Y., Liu, J., Gey van Pittius, N. C., Pym, A. S., Rubin, E. J., Sherman, D. R., Cole, S. T., and Brosch, R. (2009) Systematic genetic nomenclature for type VII secretion systems. PLoS Pathog. 5, e1000507

3. Houben, D., Demangel, C., van Ingen, J., Perez, J., Baldeón, L., Abdallah, A. M., Caleechurn, L., Bottai, D., van Zon, M., de Punder, K., van der Laan, T., Kant, A., Bossers-de Vries, R., Willemsen, P., Bitter, W., van Soolingen, D., Brosch, R., van der Wel, N., and Peters, P. J. (2012) ESX-1-mediated translocation to the cytosol controls virulence of mycobacteria. Cell. Microbiol. 14, 1287–98

4. Hsu, T., Hingley-Wilson, S. M., Chen, B., Chen, M., Dai, A. Z., Morin, P. M., Marks, C. B., Padiyar, J., Goulding, C., Gingery, M., Eisenberg, D., Russell, R. G., Derrick, S. C., Collins, F. M., Morris, S. L., King, C. H., and Jacobs, W. R. (2003) The primary mechanism of attenuation of bacillus Calmette-Guerin is a loss of secreted lytic function required for invasion of lung interstitial tissue. Proc. Natl. Acad. Sci. U. S. A. 100, 12420–5

5. Gröschel, M. I., Sayes, F., Simeone, R., Majlessi, L., and Brosch, R. (2016) ESX secretion systems: mycobacterial evolution to counter host immunity. Nat. Rev. Microbiol. 14, 677–691

6. Xu, J., Laine, O., Masciocchi, M., Manoranjan, J., Smith, J., Du, S. J., Edwards, N., Zhu, X., Fenselau, C., and Gao, L. Y. (2007) A unique Mycobacterium ESX-1 protein co-secretes with CFP-10/ESAT-6 and is necessary for inhibiting phagosome maturation. Mol. Microbiol. 66, 787–800

7. Ates, L. S., Ummels, R., Commandeur, S., van der Weerd, R., Sparrius, M., Weerdenburg, E., Alber, M., Kalscheuer, R., Piersma, S. R., Abdallah, A. M., Abd El Ghany, M., Abdel-Haleem, A. M., Pain, A., Jiménez, C. R., Bitter, W., and Houben, E. N. G. (2015) Essential role of the ESX-5 secretion system in outer membrane permeability of pathogenic mycobacteria. PLoS Genet. 11, e1005190

8. Siegrist, M. S., Unnikrishnan, M., McConnell, M. J., Borowsky, M., Cheng, T.-Y., Siddiqi, N., Fortune, S. M., Moody, D. B., and Rubin, E. J. (2009) Mycobacterial Esx-3 is required for mycobactin-mediated iron acquisition. Proc. Natl. Acad. Sci. U. S. A. 106, 18792–7

9. Tufariello, J. M., Chapman, J. R., Kerantzas, C. A., Wong, K.-W., Vilchèze, C., Jones, C. M., Cole, L. E., Tinaztepe, E., Thompson, V., Fenyö, D., Niederweis, M., Ueberheide, B., Philips, J. A., and Jacobs, W. R. (2016) Separable roles for *Mycobacterium tuberculosis* ESX-3 effectors in iron acquisition and virulence. Proc. Natl. Acad. Sci. U. S. A. 113, E348–57

10. Tinaztepe, E., Wei, J. R., Raynowska, J., Portal-Celhay, C., Thompson, V., and Philipsa, J. A. (2016) Role of metal-dependent regulation of ESX-3 secretion in intracellular survival of *Mycobacterium tuberculosis*. Infect. Immun. 84, 2255–2263

11. Garces, A., Atmakuri, K., Chase, M. R., Woodworth, J. S., Krastins, B., Rothchild, A. C., Ramsdell, T. L., Lopez, M. F., Behar, S. M., Sarracino, D. A., and Fortune, S. M. (2010) EspA acts as a critical mediator of ESX1-dependent virulence in *Mycobacterium tuberculosis* by affecting bacterial cell wall integrity. PLoS Pathog. 6, e1000957

12. Smith, J., Manoranjan, J., Pan, M., Bohsali, A., Xu, J., Liu, J., McDonald, K. L., Szyk, A., LaRonde-LeBlanc, N., and Gao, L. Y. (2008) Evidence for pore formation in host cell membranes by ESX-1-secreted ESAT-6 and its role in *Mycobacterium marinum* escape from the vacuole. Infect. Immun. 76, 5478–5487

13. De Jonge, M. I., Pehau-Arnaudet, G., Fretz, M. M., Romain, F., Bottai, D., Brodin, P., Honoré, N., Marchal, G., Jiskoot, W., England, P., Cole, S. T., and Brosch, R. (2007) ESAT-6 from *Mycobacterium tuberculosis* dissociates from its putative chaperone CFP-10 under acidic conditions and exhibits membrane-lysing activity. J. Bacteriol. 189, 6028–6034

14. Simeone, R., Bobard, A., Lippmann, J., Bitter, W., Majlessi, L., Brosch, R., and Enninga, J. (2012) Phagosomal rupture by *Mycobacterium tuberculosis* results in toxicity and host cell death. PLoS Pathog. 8, e1002507

15. van der Wel, N., Hava, D., Houben, D., Fluitsma, D., van Zon, M., Pierson, J., Brenner, M., and Peters, P. J. (2007) *M. tuberculosis* and *M. leprae* translocate from the phagolysosome to the cytosol in myeloid cells. Cell 129, 1287–1298

16. Simeone, R., Sayes, F., Song, O., Gröschel, M. I., Brodin, P., Brosch, R., and Majlessi, L. (2015) Cytosolic access of *Mycobacterium tuberculosis*: critical impact of phagosomal acidification control and demonstration of occurrence *in vivo*. PLoS Pathog. 11, e1004650

17. Abdallah, A. M., Savage, N. D. L., van Zon, M., Wilson, L., Vandenbroucke-Grauls, C. M. J. E., van der Wel, N. N., Ottenhoff, T. H. M., and Bitter, W. (2008) The ESX-5 secretion system of *Mycobacterium marinum* modulates the macrophage response. J. Immunol. 181, 7166–7175

18. Weerdenburg, E. M., Abdallah, A. M., Mitra, S., de Punder, K., van der Wel, N. N., Bird, S., Appelmelk, B. J., Bitter, W., and van der Sar, A. M. (2012) ESX-5-deficient *Mycobacterium marinum* is hypervirulent in adult zebrafish. Cell. Microbiol. 14, 728–39

19. Ates, L. S., Houben, E. N. G., and Bitter, W. (2016) Type VII secretion: a highly versatile secretion system. Microbiol. Spectr. 4, VMBF0011–2015

20. Solomonson, M., Setiaputra, D., Makepeace, K. A. T., Lameignere, E., Petrotchenko, E. V., Conrady, D. G., Bergeron, J. R., Vuckovic, M., Dimaio, F., Borchers, C. H., Yip, C. K., and Strynadka, N. C. J. (2015) Structure of EspB from the ESX-1 type VII secretion system and insights into its export mechanism. Structure 23, 571–583

21. Korotkova, N., Freire, D., Phan, T. H., Ummels, R., Creekmore, C. C., Evans, T. J., Wilmanns, M., Bitter, W., Parret, A. H. A., Houben, E. N. G., and Korotkov, K. V (2014) Structure of the *Mycobacterium tuberculosis* type VII secretion system chaperone EspG5 in complex with PE25-PPE41 dimer. Mol. Microbiol. 94, 367–382

22. Ekiert, D. C., and Cox, J. S. (2014) Structure of a PE-PPE-EspG complex from *Mycobacterium tuberculosis* reveals molecular specificity of ESX protein secretion. Proc. Natl. Acad. Sci. U. S. A. 111, 14758–63

23. Korotkova, N., Piton, J., Wagner, J. M., Boy-Röttger, S., Japaridze, A., Evans, T. J., Cole, S. T., Pojer, F., and Korotkov, K. V (2015) Structure of EspB, a secreted substrate of the ESX-1 secretion system of Mycobacterium tuberculosis. J. Struct. Biol. 191, 236–44

24. Renshaw, P. S., Lightbody, K. L., Veverka, V., Muskett, F. W., Kelly, G., Frenkiel, T. A., Gordon, S. V, Hewinson, R. G., Burke, B., Norman, J., Williamson, R. a, and Carr, M. D. (2005) Structure and function of the complex formed by the tuberculosis virulence factors CFP-10 and ESAT-6. EMBO J. 24, 2491–8

25. Ilghari, D., Lightbody, K. L., Veverka, V., Waters, L. C., Muskett, F. W., Renshaw, P. S., and Carr, M. D. (2011) Solution structure of the *Mycobacterium tuberculosis* EsxG·EsxH complex: functional implications and comparisons with other *M. tuberculosis* Esx family complexes. J. Biol. Chem. 286, 29993–30002

26. Chen, X., Cheng, H. F., Zhou, J., Chan, C. Y., Lau, K. F., Tsui, S. K. W., and Au, S. W. ngor (2017) Structural basis of the PE–PPE protein interaction in *Mycobacterium tuberculosis*. J. Biol. Chem. 292, 16880–16890

27. Daleke, M. H., Ummels, R., Bawono, P., Heringa, J., and Vandenbroucke-grauls, C. M. J. E. (2012) General secretion signal for the mycobacterial type VII secretion pathway. Proc. Natl. Acad. Sci. 109, 11342–11347

28. Patricia A. DiGiuseppe Champion, Stanley, S. A., Champion, M. M., Brown, E. J., and Cox, J. (2006) C-Terminal Signal Sequence Promotes Virulence Factor Secretion in *Mycobacterium tuberculosis*. Science. 313, 1632–1637

29. Daleke, M. H., Woude, A. D. Van Der, Parret, A. H. A., Ummels, R., Groot, A. M. De, Watson, D., Piersma, S. R., Jiménez, C. R., Luirink, J., Bitter, W., and Houben, E. N. G. (2012) Specific Chaperones for the Type VII Protein Secretion. J. Biol. Chem. 287, 31939–31947

30. Phan, T. H., Ummels, R., Bitter, W., and Houben, E. N. G. (2017) Identification of a substrate domain that determines system specificity in mycobacterial type VII secretion systems. Sci. Rep. 7, 42704

31. Abdallah, A. M., Verboom, T., Hannes, F., Safi, M., Strong, M., Eisenberg, D., Musters, R. J. P., Vandenbroucke-Grauls, C. M. J. E., Appelmelk, B. J., Luirink, J., and Bitter, W. (2006) A specific secretion system mediates PPE41 transport in pathogenic mycobacteria. Mol. Microbiol. 62, 667–79

32. Fortune, S. M., Jaeger, A., Sarracino, D. A., Chase, M. R., Sassetti, C. M., Sherman, D. R., Bloom, B. R., and Rubin, E. J. (2005) Mutually dependent secretion of proteins required for mycobacterial virulence. Proc. Natl. Acad. Sci. U. S. A. 102, 10676–81

33. Brodin, P., Majlessi, L., Marsollier, L., Jonge, M. I. de, Bottai, D., Demangel, C., Hinds, J., Neyrolles, O., Butcher, P. D., Leclerc, C., Cole, S. T., and Brosch, R. (2006) Dissection of ESAT-6 System 1 of *Mycobacterium tuberculosis* and impact on immunogenicity and virulence. Infect. Immun. 74, 88–98

34. Abdallah, A. M., Verboom, T., Weerdenburg, E. M., Gey van Pittius, N. C., Mahasha, P. W., Jiménez, C., Parra, M., Cadieux, N., Brennan, M. J., Appelmelk, B. J., and Bitter, W. (2009) PPE and PE_PGRS proteins of *Mycobacterium marinum* are transported via the type VII secretion system ESX-5. Mol. Microbiol. 73, 329–340

35. Houben, E. N. G., Bestebroer, J., Ummels, R., Wilson, L., Piersma, S. R., Jiménez, C. R., Ottenhoff, T. H. M., Luirink, J., and Bitter, W. (2012) Composition of the type VII secretion system membrane complex. Mol. Microbiol. 86, 472–484

36. Beckham, K. S. H., Ciccarelli, L., Bunduc, C. M., Mertens, H. D. T., Ummels, R., Lugmayr, W., Mayr, J., Rettel, M., Savitski, M. M., Svergun, D. I., Bitter, W., Wilmanns, M., Marlovits, C., Parret, A. H. A., and Houben, E. N. G. (2017) Structure of the mycobacterial ESX-5 type VII secretion system membrane complex by single particle analysis. Nat. Microbiol. 2, 1–8

37. Daleke, M. H., Van Der Woude, A. D., Parret, A. H. A., Ummels, R., De Groot, A. M., Watson, D., Piersma, S. R., Jiménez, C. R., Luirink, J., Bitter, W., and Houben, E. N. G. (2012) Specific chaperones for the type VII protein secretion pathway. J. Biol. Chem. 287, 31939–31947

38. Phan, T. (2018) Characterization of ESX-1 components EccA1, EspG1 and EspH reveal pivotal role of Esp substrates in the *Mycobacterium marinum* infection cycle. PLoS Pathog. 14, e1007247

39. Stanley, S. A., Raghavan, S., Hwang, W. W., and Cox, J. S. (2003) Acute infection and macrophage subversion by *Mycobacterium tuberculosis* require a specialized secretion system. Proc. Natl. Acad. Sci. U. S. A. 100, 13001–6

40. Rosenberg, O. S., Dovala, D., Stroud, R. M., Cox, J. S., Rosenberg, O. S., Dovala, D., Li, X., Connolly, L., Bendebury, A., and Finer-moore, J. (2015) Substrates control multimerization and activation of the multi-domain ATPase motor of type VII secretion. Cell 161, 501–512

41. Poulsen, C., Panjikar, S., Holton, S. J., Wilmanns, M., and Song, Y.-H. (2014) WXG100 protein superfamily consists of three subfamilies and exhibits an α-helical C-terminal conserved residue pattern. PLoS One 9, e89313

42. Gey van Pittius, N. C., Sampson, S. L., Lee, H., Kim, Y., van Helden, P. D., and Warren, R. M. (2006) Evolution and expansion of the *Mycobacterium tuberculosis* PE and PPE multigene families and their association with the duplication of the ESAT-6 (esx) gene cluster regions. BMC Evol. Biol. 6, 95

43. Houben, E. N. G., Korotkov, K. V, and Bitter, W. (2013) Take five - Type VII secretion systems of Mycobacteria. Biochim. Biophys. acta. 1844, 1707–1716

44. Unnikrishnan, M., Constantinidou, C., Palmer, T., and Pallen, M. J. (2017) The enigmatic Esx proteins: looking beyond Mycobacteria. Trends Microbiol. 25, 192–204

45. Guinn, K. M., Hickey, M. J., Mathur, S. K., Zakel, K. L., Grotzke, J. E., Lewinsohn, D. M., Smith, S., and Sherman, D. R. (2004) Individual RD1-region genes are required for export of ESAT-6/CFP-10 and for virulence of *Mycobacterium tuberculosis*. Mol. Microbiol. 51, 359–370

46. Gao, L.-Y., Guo, S., McLaughlin, B., Morisaki, H., Engel, J. N., and Brown, E. J. (2004) A mycobacterial virulence gene cluster extending RD1 is required for cytolysis, bacterial spreading and ESAT-6 secretion. Mol. Microbiol. 53, 1677–93

47. DiGiuseppe Champion, P. A., Champion, M. M., Manzanillo, P., and Cox, J. S. (2009) ESX-1 secreted virulence factors are recognized by multiple cytosolic AAA ATPases in mycobactria. Mol. Microbiol. 73, 950–962

48. Bosserman, R. E., Thompson, C. R., Nicholson, K. R., and Patricia, A. (2018) Esx paralogs are functionally equivalent to ESX-1 proteins but are dispensable for virulence in M. marinum. J. Bacteriol. 200, e00726–17

49. Harboe, M., Malin, A. S., Dockrell, H. S., Wiker, H. G., Ulvund, G., Holm, A., Jørgensen, M. C., and Andersen, P. (1998) B-cell epitopes and quantification of the ESAT-6 protein of *Mycobacterium tuberculosis*. Infect. Immun. 66, 717–23

