## Supplemental tables for "Rerouting of an Esx substrate pair from the ESX-1 type VII secretion system to ESX-5 by modifying a PE/PPE substrate pair"

Running title: PE/PPE determine system-specificity of Esx substrates

[*]To whom correspondence should be addressed: Edith N.G. Houben: Vrije Universiteit Amsterdam, Section Molecular Microbiology, de Boelelaan 1108, 1081 HZ Amsterdam, the Netherlands. Tel.: +31 20 5983579;

#These authors contributed equally to this work.

**Material included:**

- Table S1
- Table S2
- Table S3

**Table S1.** Strains used in this study.

| Strain | References |
|---|---|
| WT *M. marinum* M | (29) |
| *eccCb₁* mutant (ESX-1 mutant) | (32) |
| $eccC_5$ mutant (ESX-5 mutant) | This study, complete deletion of $eccC_5$ in the genome of *M. marinum* M background strain |

**Table S2.** Plasmids used in this study.

| Plasmids | Characteristics | Gene origin | Reference |
|---|---|---|---|
| pSMT3::*esxB_1/EsxA_1 WT.HA* | hsp60 promoter, hygroR | *M. marinum* M | This study |
| pMV::*pe35/ppe68_1 WT.FLAG* | hsp60 promoter, KanR, integrative | *M. marinum* M | This study |
| pSMT3::*pe35/ppe68_1-FLAG /esxB_1/esxA_1.HA WT* | hsp60 promoter, hygroR, all four genes are WT | *M. marinum* M | This study |
| pSMT3::*pe35/ppe68_1-FLAG /esxB_1/esxA_1.HA* SINGLE SWAP | hsp60 promoter, hygroR, *ppe68_1* carried *espG*$_5$ chaperone binding domain of *ppe18* | *M. marinum* M | This study |
| pSMT3::*pe35/ppe68_1-FLAG /esxB_1/esxA_1.HA* DOUBLE SWAP | similar to number 4, plus *esxB_1* carries secretion signal of *esxM* | *M. marinum* M | This study |
| pSMT3::*pe35/ppe68_1-FLAG /esxB_1/esxA_1.HA* TRIPLE SWAP | similar to number 5, plus *pe35* carries secretion signal of *pe31* | *M. marinum* M | This study |

**Table S3.** Primers used in this study.

| Primers used in this study | Sequence | Purpose of use |
|---|---|---|
| pe35-NheI-Fw | CCCGCTAGCATGCGATCCATGTCTTTTGA | cloning *pe35/ppe68_1mmar* into pSMT3 |
| ppe68_Flag Rv | TCACTTGTCGTCATCGTCTTTGTAGTCCCAGT CGTCGTCGTCATC | to clone the cluster *pe35/ppe68_1/esxB_1/EsxA_1* into pSMT3 |
| Flag_esxB Fw | GACTACAAAGACGATGACGACAAGTGAGAGT CGTTGCTAAAAAGGACTTTC | to clone the cluster *pe35/ppe68_1/esxB_1/EsxA_1* into pSMT3 |
| esxA HA tag BamHI Rv | CCCCCGGATCCTTAAGCTAAGCATAATCAGG A ACATCATACGGATAGCCGAACATCCCCG | to clone the cluster *pe35/ppe68_1/esxB_1/EsxA_1* into pSMT3 |
| ppe68_1 Flag_HindIII RV | CCCCCAAGCTTTCACTTGTCGTCATCGTCTTT GT AGTCCCAGTCGTCGTCGTCATC | to clone *pe35/ppe68_1* with FLAG tag into pMV |
| pe35 EcoRI FW | CCCCCGAATTCATGCGATCCATGTCTTTTGAC C CCG | to clone *pe35/ppe68_1* with FLAG tag into pMV |
| pe35_1 SWAP pe31 ss Rv | CGTTGGCGGTTTCGGTGCTCAGGTACGAGGC C GCGATCTGCCGC | to clone the secretion signal of *pe31* into *pe35_1* |
| pe31 ss flank ppe68_1 FW | CACCGAAACCGCCAACGCCGTGGCATCTCAA T AGTCGGCCTGCCAAC | to clone the secretion signal of *pe31* into *pe35_1* |
| esxB Fw esxN | GTCCTCGCAAATGGGCTTCTGACGCGCAAAG C CAC | exchange 15 aa of secretion signal of *esxB* into *esxM* |
| esxB Rv esxM | CTAGCCGCGGAGGACCTGCTGGGAAGCCTGC T CTTGCTGCTCGTAGTTGTTACCGGCCTGACGG | exchange 15 aa of secretion signal of *esxM* into *esxB* |
| ppe68_1 Flag_HindIII RV | CCCCCAAGCTTTCACTTGTCGTCATCGTCTTT GT AGTCCCAGTCGTCGTCGTCATC | to clone *pe35/ppe68_1* with FLAG tag into pMV |
| pe35 EcoRI FW | CCCCCGAATTCATGCGATCCATGTCTTTTGAC CC CG | to clone *pe35/ppe68_1* with FLAG tag into pMV |
| esxB NheI Fw | CCCCCGCTAGCATGGCAGAGATGAAGACCGA T GC | to clone *esxB_1/esxA_1* into pSMT3 |
